# Topically delivered 22 nt siRNAs enhance RNAi silencing of endogenous genes in two species

**DOI:** 10.1101/2020.04.20.052076

**Authors:** Bill Hendrix, Wei Zheng, Matthew J. Bauer, Ericka R. Havecker, Jennifer T. Mai, Paul H. Hoffer, Rick Sanders, Brian D. Eads, Amy Caruano-Yzermans, Danielle N. Taylor, Chelly Hresko, Janette Oakes, Alberto B. Iandolino, Michael J. Bennett, Jill Deikman

## Abstract

22 nt miRNAs or siRNAs have been shown to specifically induce production of transitive (secondary) siRNAs for targeted mRNAs. An abrasion method to deliver dsRNAs into leaf cells of intact plants was used to investigate the activities of 21 and 22 nt siRNAs in silencing genes in *Nicotiana benthamiana* and *Amaranthus cruentus*. We confirmed that both 21 and 22 nt siRNAs were able to silence a green fluorescent protein (*GFP*) transgene in treated leaves of *N. benthamiana*, but systemic silencing of *GFP* occurred only when the guide strand contained 22 nt. Silencing in the treated leaves of *N. benthamiana* was demonstrated for 3 endogenous genes: *magnesium cheletase subunit I* (*CHL-I*), *magnesium cheletase subunit H* (*CHL-H*), and *GUN4*. However, systemic silencing of these endogenous genes was not observed. Very high levels of transitive siRNAs were produced for *GFP* in response to treatment with 22 nt siRNAs, but only low levels were produced in response to a 21 nt siRNA. The endogenous genes tested also had more transitive siRNAs produced in response to 22 nt siRNAs, but the response varied from weak (*CHL-I*) to strong (*CHL-H*). 22 nt siRNAs produced greater local silencing phenotypes than 21 nt siRNAs for *GFP, CHL-H* and *GUN4* in *N. benthamiana*. The special activity of 22 nt siRNAs in producing a greater local phenotype and induction of elevated levels of transitive siRNAs was also shown in *A. cruentus* for the *CHL-H* gene. These experiments suggest a functional role for transitive siRNAs in amplifying the RNAi response.

## Introduction

RNA interference (RNAi) is a gene silencing mechanism used by plants and other eukaryotes to regulate the expression of endogenous genes, to silence transposons (and other heterochromatic regions), and defend against virus infection (Baulcombe 2004). It is achieved by the action of a small RNA (sRNA) with complementarity to the target gene sequence bound to an argonaute (AGO) protein to form an RNA-induced silencing complex (RISC). Silencing can occur at the transcriptional or post-transcriptional level, depending on the specific forms of these components. RNAi has been used to create a number of valuable traits in plants, and there is great potential for future applications to crop improvement (Frizzi and Huang 2010). Recently, plants have been shown to respond to RNA applied topically to leaf surfaces (Dalakouras et al. 2016; Mitter et al. 2017; Huang et al. 2018; Niehl et al. 2018; Sammons et al. 2018; Dubrovina et al. 2019), and this delivery route would expand the opportunities for using RNAi for agronomic benefit. In addition, topical delivery of sRNAs to plant cells provides a tool that can be used to investigate details of the RNAi mechanisms in plants.

The 21-24 bp double-stranded duplexes that initiate RNA silencing in plants are produced by Dicer-like (DCL) RNAse III family enzymes acting on various precursors, including miRNA gene hairpin-structured transcripts (Yu et al. 2017) and long dsRNAs of various sources including viruses, transposons or transgenic hairpins (Axtell 2013). These sRNA duplexes contain 2 bp 3’ overhangs (Axtell 2013). In forming the RISC, the guide strand complementary to the target mRNA is selectively bound to an AGO protein, while the passenger strand is degraded (Fang and Qi 2016). For post-transcriptional silencing, the RISC interacts with the complementary mRNA and either cleaves (slices) the complementary mRNA (Arribas-Hernández et al. 2016) or acts to repress translation (Li et al. 2013; Fang and Qi 2016).

Secondary siRNAs (also referred to as transitive siRNAs) can be produced from transcripts targeted by miRNAs or siRNAs (Moissiard et al. 2007). These secondary siRNAs may amplify silencing of the primary target or act in trans to silence members of a gene family, such as the well-studied example of NBS-LRR disease-resistance mRNAs (Axtell 2013). Secondary siRNA production requires RNA-dependent RNA polymerase 6 (RDR6) and DCL4 and is sometimes phased (Axtell 2013). While there are multiple factors that can initiate secondary siRNA production (Axtell 2013), they are produced in response to 22 nt miRNAs, while 21 nt miRNAs do not usually have this effect (Chen et al. 2010; Cuperus et al. 2010). The functional significance of 22 nt miRNAs was shown using artificial miRNAs (amiRNAs) in transgenic Arabidopsis plants, where whole-plant silencing of a targeted chalcone synthase (CHS) gene was only achieved with a 22 nt amiRNA, and not with the corresponding 21 nt amiRNA (McHale et al. 2013). This effect was dependent on RDR6, and the generation of phasiRNAs was demonstrated. The authors hypothesized that the enhanced silencing could be due to greater mobility of siRNAs (Tretter et al. 2008; Felippes et al. 2011) or to production of a significant amount of additional siRNAs homologous to CHS allowing gene knock-down throughout the plant.

There are several pathways for the generation of 22 nt siRNAs or miRNAs in plant cells. DCL2 processes 22 nt siRNAs from long dsRNAs, and its activity is critical for both the production of secondary siRNAs and the transitive silencing of transgenes (Mlotshwa et al. 2008; Parent et al. 2014; Taochy et al. 2017; Chen et al. 2018). Alternatively, 22 nt miRNAs can be generated by processing by DCL1 of an asymmetric precursor miRNA to generate a 21/22 nt miRNA (Manavella et al. 2012), by imprecise cutting by DCL1 (Li et al. 2016), or by mono-uridylation by HEN1 Supressor1 (HESO1) of a 21 nt miRNA (Fei et al. 2018). The exact structure of a miRNA or siRNA that can promote production of secondary siRNAs is unclear, and there are instances where 21 nt miRNAs can trigger production of secondary siRNAs (Manavella et al. 2012). Additionally, examples were shown where secondary siRNAs can be produced when the passenger strand of the miRNA duplex is 22 nt but the miRNA is 21 nt (Manavella et al. 2012). These researchers proposed that the critical structural feature for induction of secondary siRNAs is not the specific size of the miRNA, but asymmetry in the miRNA duplex structure. Recent work showing formation of a 22 nt miRNA that is associated with secondary siRNA production by mono-uridylation of a 21 nt miRNA (Fei et al. 2018) would support the importance of miRNA length alone, although it doesn’t rule out other mechanisms.

Silencing by RNAi can spread from a site of initiation to distal parts of a plant. This phenomenon has been observed in multiple species using grafting of genotypes containing silencing transgenes, by initiation of local silencing using *Agrobacterium* infiltration of hairpin constructs, by expression of hairpins or amiRNAs under control of tissue specific promoters, or by topical introduction of a dsRNA (Palauqui et al. 1997; Voinnet and Baulcombe 1997; McHale et al. 2013; Dalakouras et al. 2016). The signal for systemic silencing has not been conclusively determined but is generally thought to be an RNA (Melnyk et al. 2011; Taochy et al. 2017; Zhang et al. 2019). sRNAs can move from cell-to-cell in plants through plasmodesmata or longer distances through vascular tissues (Melnyk et al. 2011; Liu and Chen 2018). Cell-to-cell movement combined with template-dependent signal amplification is another mechanism described for long-distance sRNA movement (Liang et al. 2012). Recent work has highlighted the critical role of *DCL2*, and thus 22 nt siRNAs, in systemic RNAi (Taochy et al. 2017; Chen et al. 2018).

We used an abrasion-based delivery method to introduce dsRNAs into leaf cells of intact plants for two plant species and confirmed enhanced activity of 22 nt siRNAs compared to 21 nt siRNAs. 22 nt siRNAs induced systemic silencing of a *GFP* transgene in *N. benthamiana* with this delivery method, while 21 nt siRNAs did not. However, we did not observe systemic silencing for any of the 3 endogenous genes targeted. 22 nt siRNAs produced greater local phenotypes than 21 nt siRNAs for 3 of the 4 gene targets tested. 22 nt siRNAs induced secondary siRNAs production for all target mRNAs, but to different levels. Very high levels of secondary siRNAs were produced for the *magnesium cheletase H* (*CHL-H*) gene, but systemic silencing was still not observed. This low-tech RNA delivery method is simple to use and has utility for future study of factors required for efficient RNAi in plants.

## Results

### *GFP* transgene silencing in *N. benthamiana* by topically applied dsRNA

The *GFP* transgene was silenced using a sandpaper abrasion method (Huang et al. 2018) to introduce dsRNAs into leaf cells of *N. benthamiana* line 16C (Ruiz et al. 1998). An efficacious dsRNA to silence *GFP* was selected by calculating a Reynolds score for all possible 19 mers (Reynolds et al. 2004), and then testing high scoring sequences as 22 nt siRNAs with 2 nt 3’ overhangs in protoplasts prepared from tobacco BY-2 cells expressing a *GFP* transgene. The 22 nt siRNA showing the greatest silencing in protoplasts was used as the base sequence for studies using topical dsRNA treatment of intact plants (Table 1).

**Table 1.**
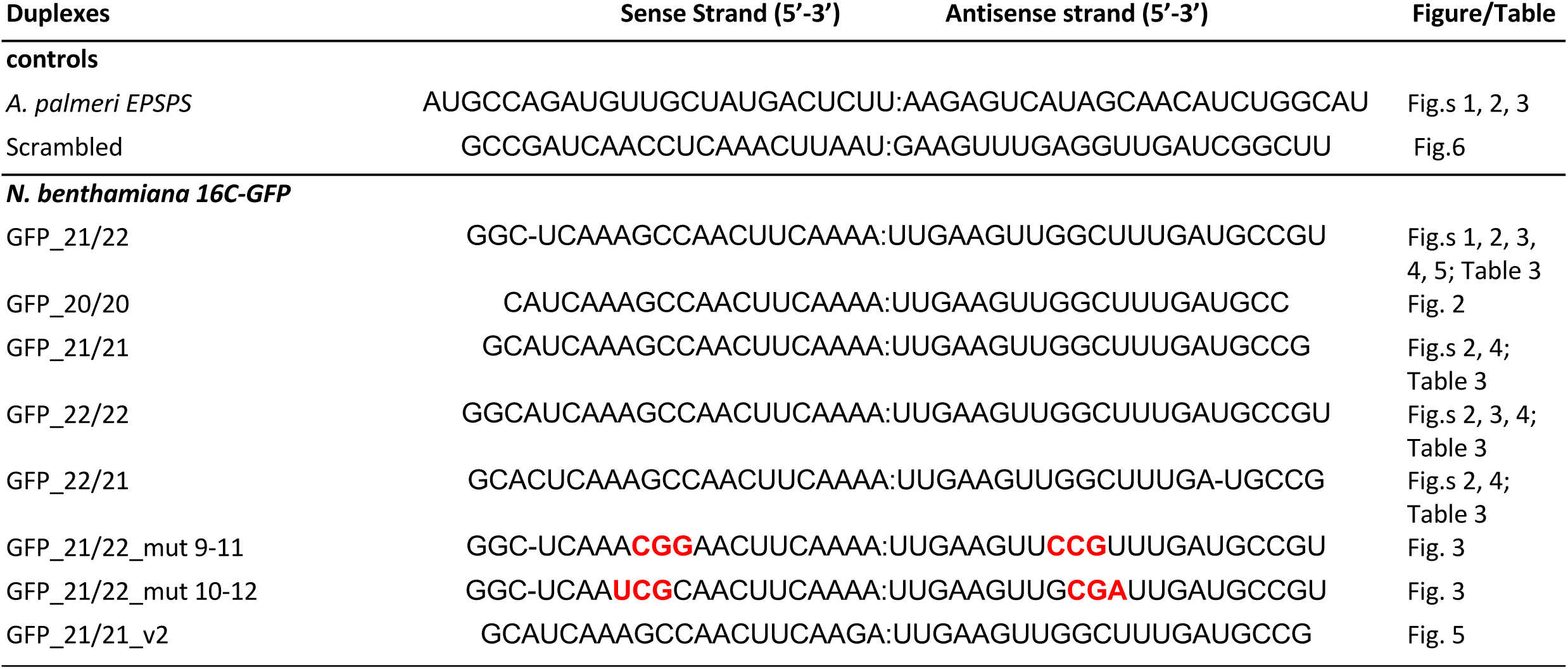
dsRNAs targeting GFP or a control sequence. Letters in red indicate introduced mutations.

GFP fluoresces green under UV light, and GFP gene suppression can be detected by the unmasking of red fluorescence emitted from chlorophyll. We observed that *GFP* silencing was visible under UV light on treated leaves as soon as 2 days after treatment (dat), and phenotype on treated leaves was maximal by 8 days after treatment (Fig. 1A-F). Leaves treated with a control dsRNA (24 bp of the 5-enolpyruvylshikimate-3-phosphate synthase (EPSPS) coding sequence from *Amaranthus palmeri*) did not show development of red fluorescent spots (Fig. 1B). Systemic silencing in veins of upper leaves was strong at 13 days after treatment in this experiment and continued to progress during the life of the plant. Quantification of an RNA gel blot showed that expression of the *GFP* transgene in treated leaves was reduced 25% 1 day after treatment and 67% 2 days after treatment (Fig. 1G).

**Figure 1.**
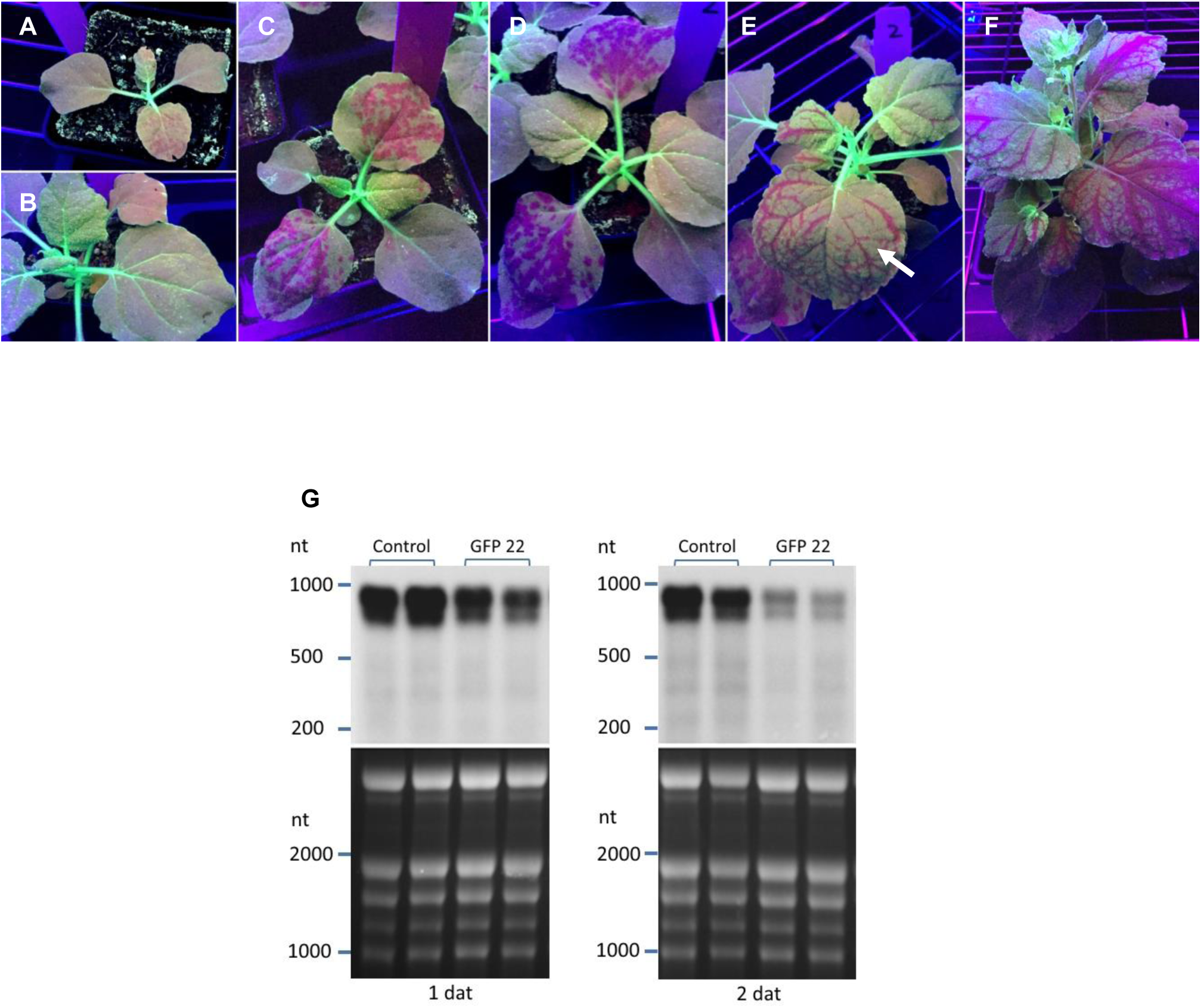
Development of *GFP* silencing over time in response to topically-applied dsRNA. *GFP* silencing appeared as red spots on treated leaves when plants were viewed under UV light as soon as 2 days after treatment (dat) and systemic silencing was visible in veins of upper leaves (arrow) by 13 days after treatment. dsRNA was applied onto leaves 3 and 4 of 14-day old plants followed by abrasion with #600 grit sandpaper. Plants were treated with GFP 21/22 siRNA (A and C-F) or a control RNA (B, 24 bp dsRNA from *A. palmeri EPSPS*), and photos were taken under UV lights at the following time points: A, 2 dat; B and C, 5 dat; D, 8 dat; E, 13 dat; F, 26 dat. G, RNA gel blot probed with GFP sequence (upper panels) and gel stained with ethidium bromide (lower panels). Samples from treated leaves were collected 1 and 2 days after treatment.

To better understand the sRNA structural requirements for optimal activity, we treated 16-day old N. benthamiana leaves 3, 4 and 5 (counted from the base of the plant) with siRNAs targeting GFP from 20 to 22 nt, including hybrid molecules containing one strand with 22 nt and the other with 21 nt (Table 1). Plants were examined at 6 dat under blue light to observe GFP silencing phenotypes (Fig. 2). Note that the treated leaves are circled, and red spots indicate silencing of the *GFP* transgene. We consistently observed that leaves 1 and 2 appeared red without siRNA treatment and hypothesize that the *GFP* transgene is not strongly expressed in these tissues at this developmental stage. However, untreated leaves 3 and older consistently showed green GFP fluorescence, and silencing of *GFP* could be observed by appearance of red spots in these leaves. While some silencing was evident at 6 dat from all GFP siRNAs tested from 20 nt to 22 nt, silencing in treated leaves was strongest with the 22/22 or 21/22 configurations, which both included the 22 nt sequence in the guide strand of the dsRNA. Systemic silencing was observed at 15 dat in plants that had been treated with siRNAs containing a 22 nt guide strand, and both 22/22 and 21/22 siRNA configurations produced systemic silencing (Fig 2). A sequence containing 22 nt in the passenger strand with a 21 nt guide strand (22/21) did not induce systemic silencing (Fig 2).

**Figure 2.**
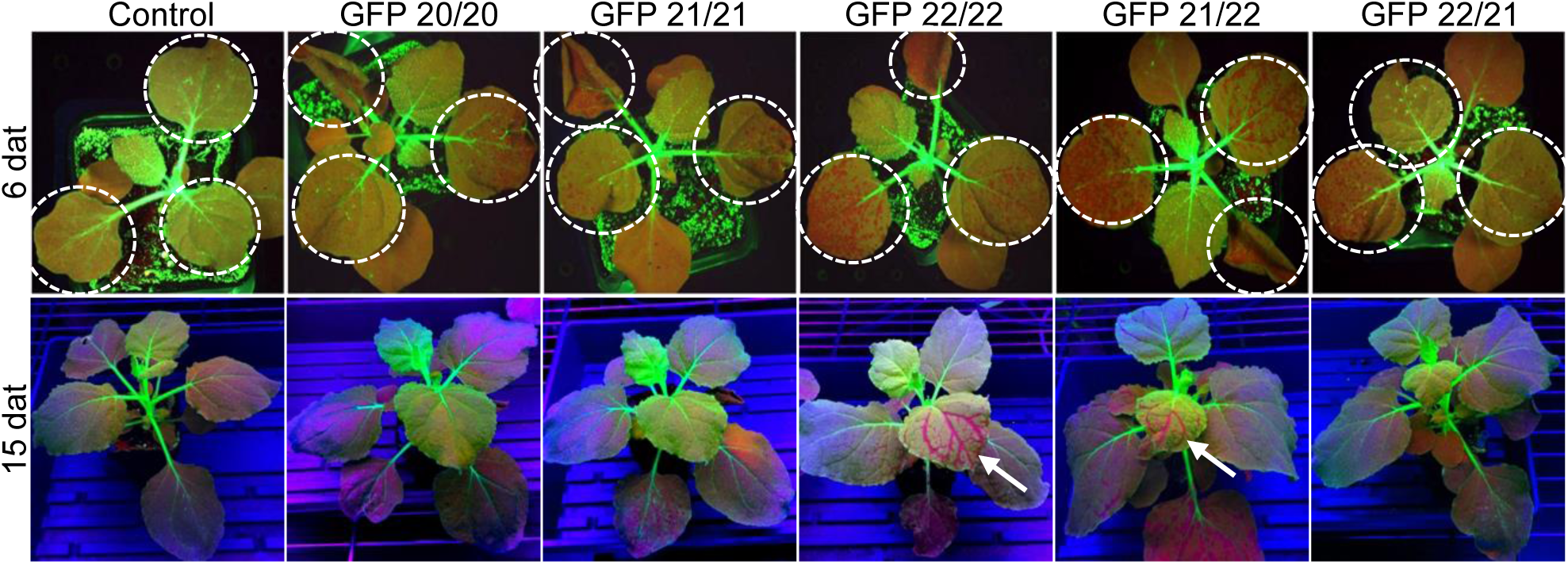
An siRNA with 22 nt in the guide strand was required for systemic *GFP* silencing in *N. benthamiana*. dsRNA (25 μl dsRNA at 1 μg/μl in 0.01% Silwet L77) was applied onto true leaves 3, 4, and 5 of 16-day old plants followed by abrasion with #600 grit sandpaper (3 plants per treatment). Plants were photographed under strong blue lights at 6 dat and UV lights at 15 dat. One representative plant per treatment is shown and treated leaves are circled. Arrows point to leaves showing systemic *GFP* silencing. Control, 24 bp dsRNA from *A. palmeri EPSPS*.

To further confirm *GFP* silencing observed with the abrasion method was a result of RNAi, siRNAs containing mutations of the predicted slice site were tested (Table 1). Consistent with topically-delivered siRNAs acting in the RNAi pathway, mutations of the GFP siRNAs from 9-11 (GGC to CCG) or 10-12 (GCU to CGA) nt, which would alter the AGO slice site, greatly decreased silencing activity (Fig. 3A).

**Figure 3.**
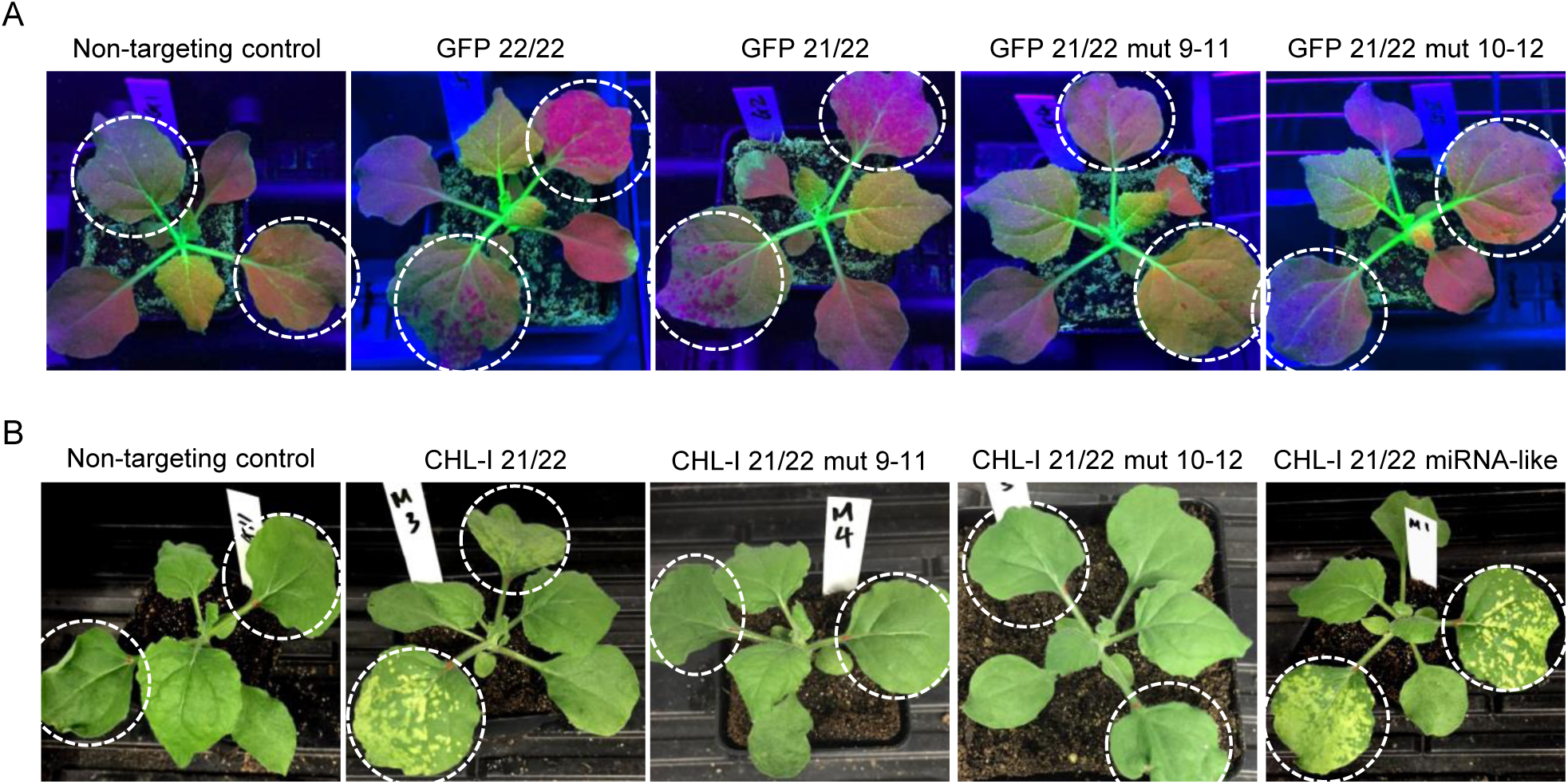
Activity of siRNAs with slice site mutations was consistent with an RNAi mode-of-action. dsRNA (20 μl dsRNA at 1 μg/μl in 0.01% Silwet L77) was applied and evenly spread onto the leaves 3 and 4 of 14-day plants followed by abrasion with #600 grit sandpaper (2 plants per treatment). The treated leaves are circled. A, A representative plant treated with *GFP*-targeting or control dsRNAs. The plants were photographed under UV lights at 5 dat. B, A representative plant treated with *CHL-I*-targeting or control dsRNAs. The plants were photographed under white LED lights at 5 dat. Control, 24 bp dsRNA from *A. palmeri EPSPS*.

### Silencing of the endogenous *magnesium cheletase I (CHL-I)* gene in *N. benthamiana*

We then targeted an endogenous gene in *N. benthamiana* encoding *magnesium cheletase I (CHL-I)*, which is involved in chlorophyll biosynthesis (Kruse et al. 1997). Several different CHL-I 21/22 siRNAs were tested with sandpaper delivery in *N. benthamiana* line 16C, and the most efficacious sequence was used in further experiments (data not shown; Table 2, CheI_21/22). Silencing of *CHL-I* resulted in appearance of yellow spots where RNA had been delivered (Fig 3B). 22 nt variants of the 21/22 CHL-I siRNA were made containing mutations of nucleotides at positions 9-11 (UCU to AUC) or 10-12 (CUC to UCG) of the guide strand to inactivate the predicted AGO slice site. These siRNAs were not active in silencing *CHL-I* using sandpaper-based dsRNA delivery, supporting an RNAi mode-of-action for this delivery method (Fig 3B). An siRNA containing mismatches in the passenger stand (positions 9, 13, and 19) was created to mimic a miRNA structure (Table 2 and Fig 3B, “CHL-I 21/22 miRNA-like”) (Jones-Rhoades and Bartel 2004). This structure had activity comparable to the 21/22 siRNA that included complete sequence identity to the CHL-I target on both strands (Fig 3B, “CheI 21/22”). These results are all consistent with the hypothesis that silencing of *CHL-I* with topically-applied dsRNA occurs through the RNAi pathway. Plants were observed for 2 weeks after treatment with CHL-I siRNAs, but systemic silencing was never observed for this gene, although systemic silencing for the GFP transgene was strong at this time point (data not shown).

**Table 2.**
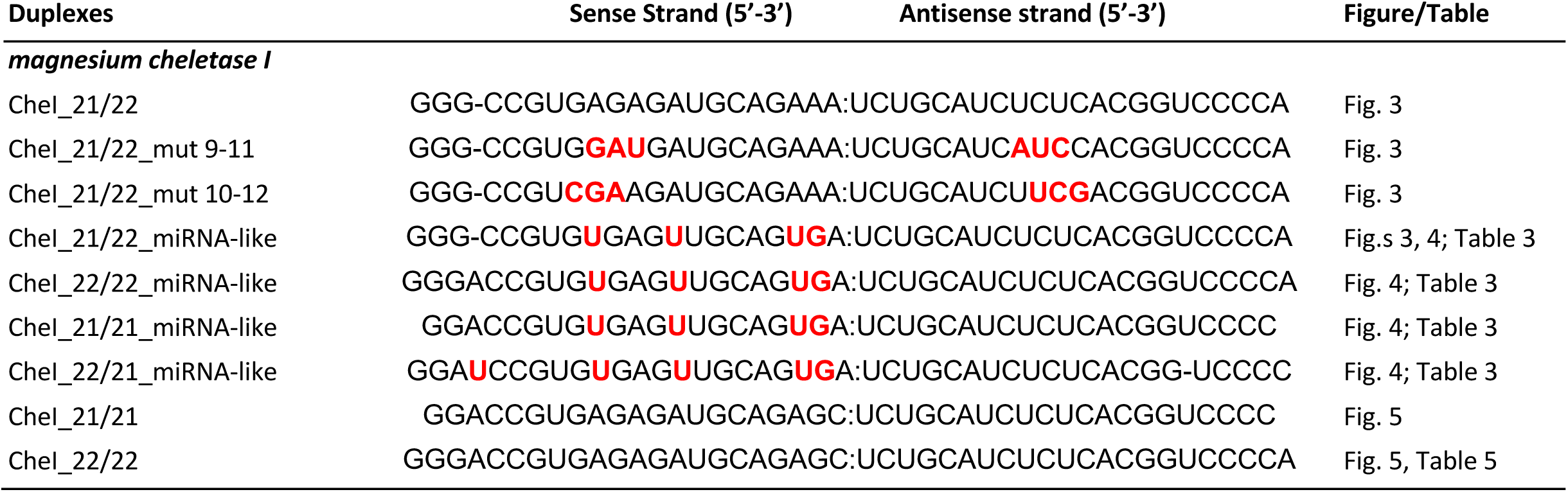
dsRNAs targeting *N. benthamiana* CHL-I gene. Red letters indicate introduced mutations or mismatches from the antisense strand sequence.

### Transitive sRNA production after topical treatment with siRNAs

Since 22 nt miRNAs have been associated with transitive sRNAs or phasiRNAs (Axtell 2013), we examined sRNAs present in tissue showing silencing phenotype for *GFP* or *CHL-I* after treatment with siRNAs containing a 21 or 22 nt sequence as either the guide or passenger strand, or both (Tables 1 and 2). The base siRNA for CHL-I was the “miRNA-like form” demonstrated to have activity comparable to the simple 21/22 siRNA (Fig. 3, Table 2), while the base GFP siRNA was a 22/22 siRNA. Three-week old plants were treated with siRNAs using sandpaper abrasion, and phenotypic tissue was harvested 4 days after treatment for sRNA analysis. Phenotype was observed but not quantified in this experiment, and all treatments with siRNAs targeting genes displayed the expected phenotypes. A baseline level of sRNAs for the *GFP* transgene and for *CHL-I* were observed in a control that involved application of buffer followed by sandpaper rolling (Table 3). Some contamination with the applied siRNA sequence was noted in this control treatment (Fig 4A and B). We concluded that this contamination occurred at a point after delivery of applied siRNAs, since it was not associated with production of transitive sRNAs. Treatment with a 22 nt siRNA in the guide strand targeting GFP (21/22 or 22/22) resulted in very high levels of transitive sRNAs, from 1000 to 1600 times background levels (Table3). sRNA levels were elevated both 5’ and 3’ to the siRNA binding site, but the amount of sRNAs was greatest 3’ of the siRNA binding site (Table 3 and Fig 4A). siRNAs targeting *GFP* containing 21 nt in the guide strand, even with 22 nt in the passenger strand (22/21 or 21/21), induced only low levels of transitive sRNA production. These levels were 14 times the background level, but the 22 nt siRNA targeting *GFP* resulted in approximately 70 times more transitive sRNAs 3’ to the siRNA binding site compared to the 21 nt siRNA. The transitive sRNAs produced in response to any siRNA were mostly 21 nt in length, but some 22 and 24 nt sRNAs were also produced (Fig 4A inset graphs). No significant differences in sRNA size ratios were associated with a particular siRNA configuration.

**Table 3.** sRNAs mapping to GFP and CHL-I genes after treatment with siRNAs with different structures.

| Target gene | sRNA Type | siRNA | Sum of normalized small RNA counts 19-25nt |  |
| --- | --- | --- | --- | --- |
|  |  |  | GFP | CHL-I |
| GFP | 5' Transitive | 21/22 | 84,701 | 1,662 |
|  | 3' Transitive |  | 449,730 | 15 |
|  | 5' Transitive | 22/22 | 58,392 | 895 |
|  | 3' Transitive |  | 294,609 | 25 |
|  | 5' Transitive | 21/21 | 4,410 | 915 |
|  | 3' Transitive |  | 3,998 | 122 |
|  | 5' Transitive | 22/21 | 6,170 | 1,444 |
|  | 3' Transitive |  | 4,148 | 62 |
| CHL-I | 5' Transitive | 21/22 | 9,542 | 1,101 |
|  | 3' Transitive |  | 253 | 7,208 |
|  | 5' Transitive | 22/22 | 2,825 | 1,336 |
|  | 3' Transitive |  | 478 | 14,175 |
|  | 5' Transitive | 21/21 | 974 | 143 |
|  | 3' Transitive |  | 115 | 14 |
|  | 5' Transitive | 22/21 | 2,618 | 4,450 |
|  | 3' Transitive |  | 252 | 39 |
| None | 5' Transitive | Sand paper only | 3,206 | 866 |
|  | 3' Transitive |  | 281 | 11 |

**Figure 4.**
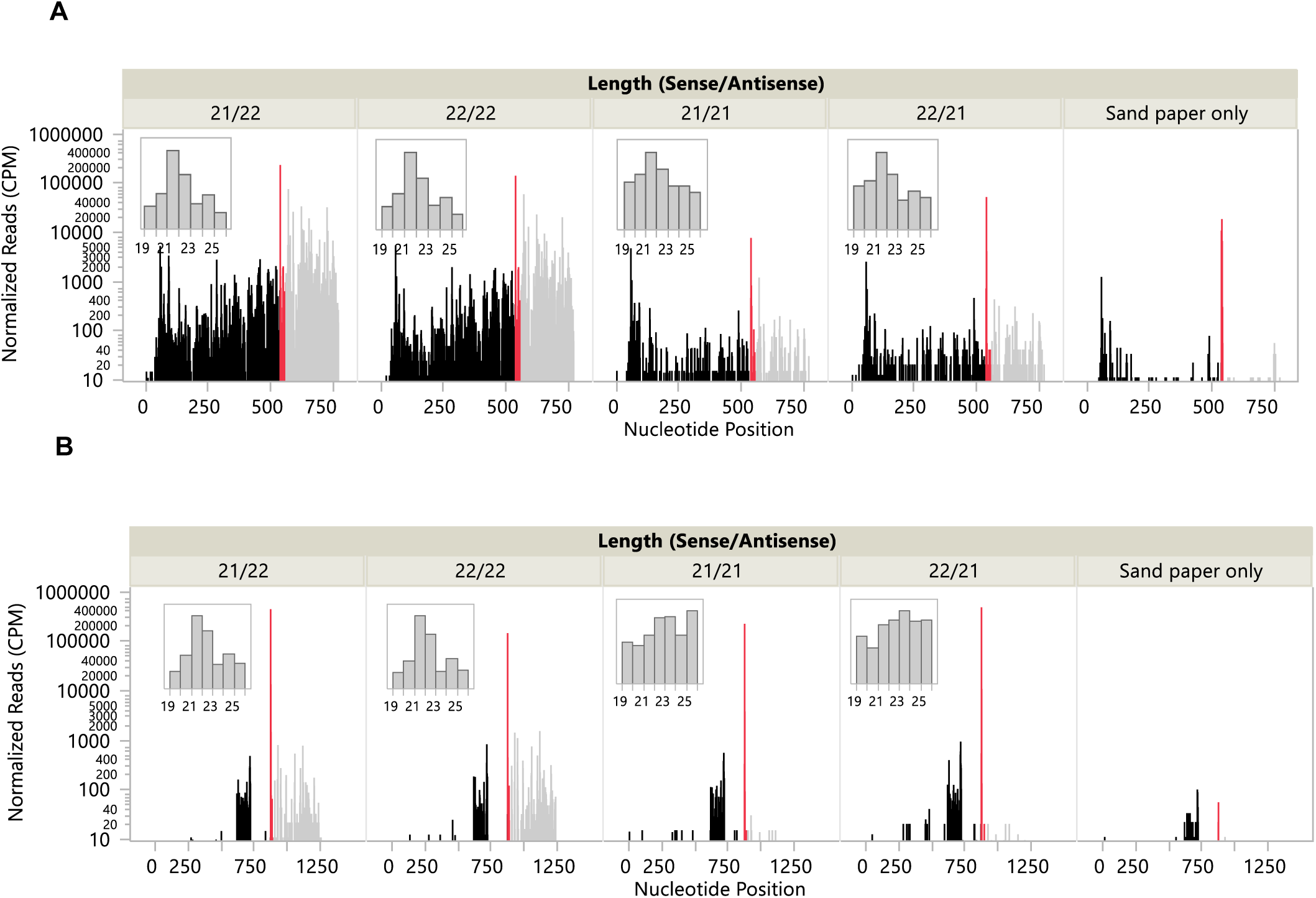
Transitive sRNA production was greater when the guide strand of the siRNA contained 22 nucleotides. siRNAs were applied to the 2 youngest partially expanded leaves of 3-week old plants (3 reps), and treated leaves were lightly abraded with sandpaper. Four days after treatment phenotypic tissue was harvested from the 2 plants with the best phenotypes and RNA was isolated and sRNA sequencing was conducted. A, sRNAs were mapped to the GFP transcript for samples from plants treated with dsRNAs targeting *GFP* or the sandpaper-only control. Insets show distribution of sRNA sizes. B, sRNAs were mapped to the CHL-I transcript for samples from plants treated with siRNAs targeting *CHL-I* or the sandpaper-only control. Insets show distribution of sRNA sizes.

The CHL-I target also produced transitive sRNAs in response to targeting 22 nt siRNAs, but the response was much weaker than for the *GFP* transgene, even though the tissue harvested had strong gene knock-down as observed by the chlorotic phenotype. For the CHL-I target, only a low level of transitive sRNAs were generated in response to treatment with siRNAs containing 22 nt in the guide strand (21/22 or 22/22), and these sRNAs were only produced 3’ to the siRNA binding site (Table 3 and Fig 4B). No increase in transitive sRNAs 3’ to the siRNA binding site were observed when the siRNA had 21 nt in the guide strand. The numbers of sRNAs produced 5’ to the siRNA binding site were more variable among samples, and no clear response to an siRNA type was observed. Comparing the response to 22 nt siRNAs, about 20 times more transitive sRNAs 3’ to the siRNA binding site were produced for GFP compared to CHL-I. The size distribution of CHL-I transitive sRNAs were similar to those produced for GFP, with 21 nt having the greatest representation, but 22 and 24 nt sRNAs were also made. This size distribution was not observed when the siRNAs had 21 nt guide strands, consistent with lack of transitive sRNA production.

### Production of higher levels of transitive sRNAs in response to 22 nt siRNAs was associated with greater silencing activity

To expand our understanding of silencing of endogenous genes by topically-applied dsRNAs, we targeted two other endogenous genes in *N. benthamiana* that when silenced produce chlorotic tissue. These genes were *magnesium cheletase-H* (*CHL-H*), another sub-unit of the magnesium cheletase enzyme (Kruse et al. 1997), and *GUN4*, a gene involved in plastid retrograde signaling (Larkin et al. 2003). While topical treatment with 22 nt siRNAs targeting these genes (Table 4) did produce yellow spots in the treated leaves, no systemic silencing was observed (data not shown).

**Table 4.**
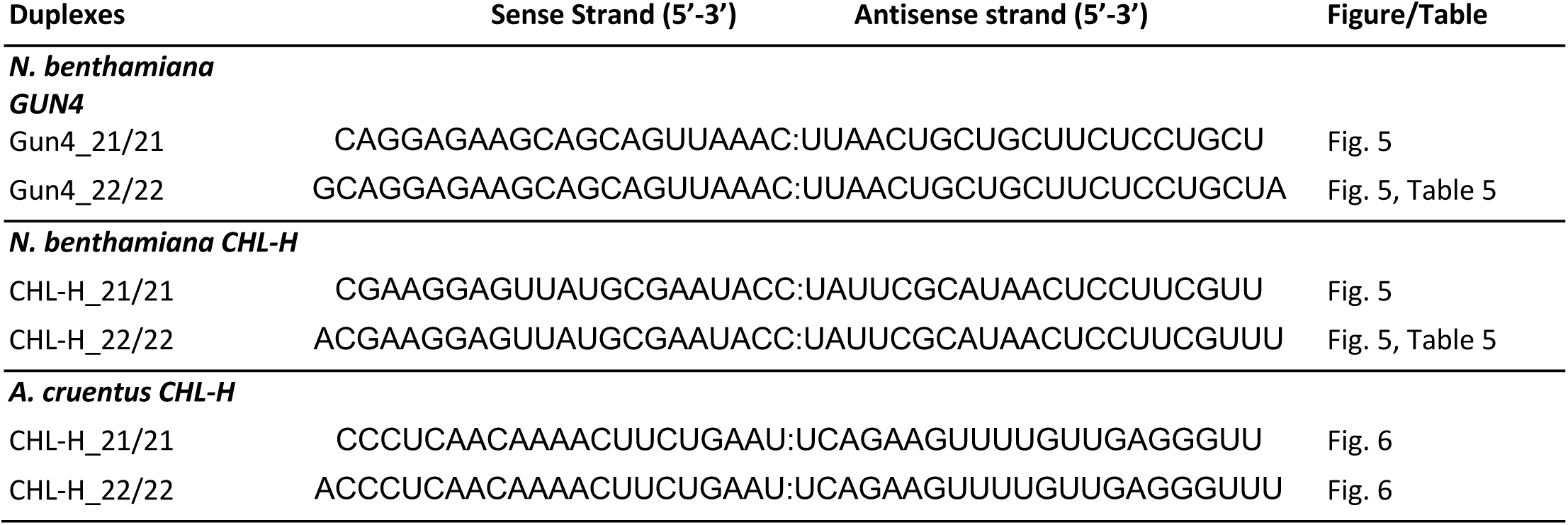
dsRNAs targeting *GUN4* and *CHL-H* genes.

An experiment was conducted to determine whether 21 vs 22 nt siRNAs had differential effects on gene silencing within the treated leaves. 21 or 22 bp siRNAs targeting *GFP, CHL-I, CHL-H* and *GUN4* were delivered to leaves of *N. benthamiana* line 16C at a dose that allowed separation of phenotypic spots. 22 nt siRNAs showed greater activity than 21 nt siRNAs in generating silencing phenotypes for *GFP, CHL-H*, and *GUN4*, and for these genes the phenotype was increased approximately 4.4-, 5.5- and 2.5-fold, respectively (Fig. 5A and B). However, the 21 nt siRNA targeting *CHL-I* produced 7.9 times greater silencing phenotype than the 22 nt siRNA. These effects were confirmed in multiple experiments. mRNA levels for the targeted endogenous genes were measured in phenotypic spots from these treatments and compared to mRNA levels from green tissue of controls. 22 bp siRNAs had greater activity than 21 bp siRNAs in reduction of mRNA levels for *GUN4* and *CHL-H* (Fig. 5C). However, 21 and 22 nt siRNAs had similar activity in reducing mRNA levels for *CHL-I*.

**Figure 5.**
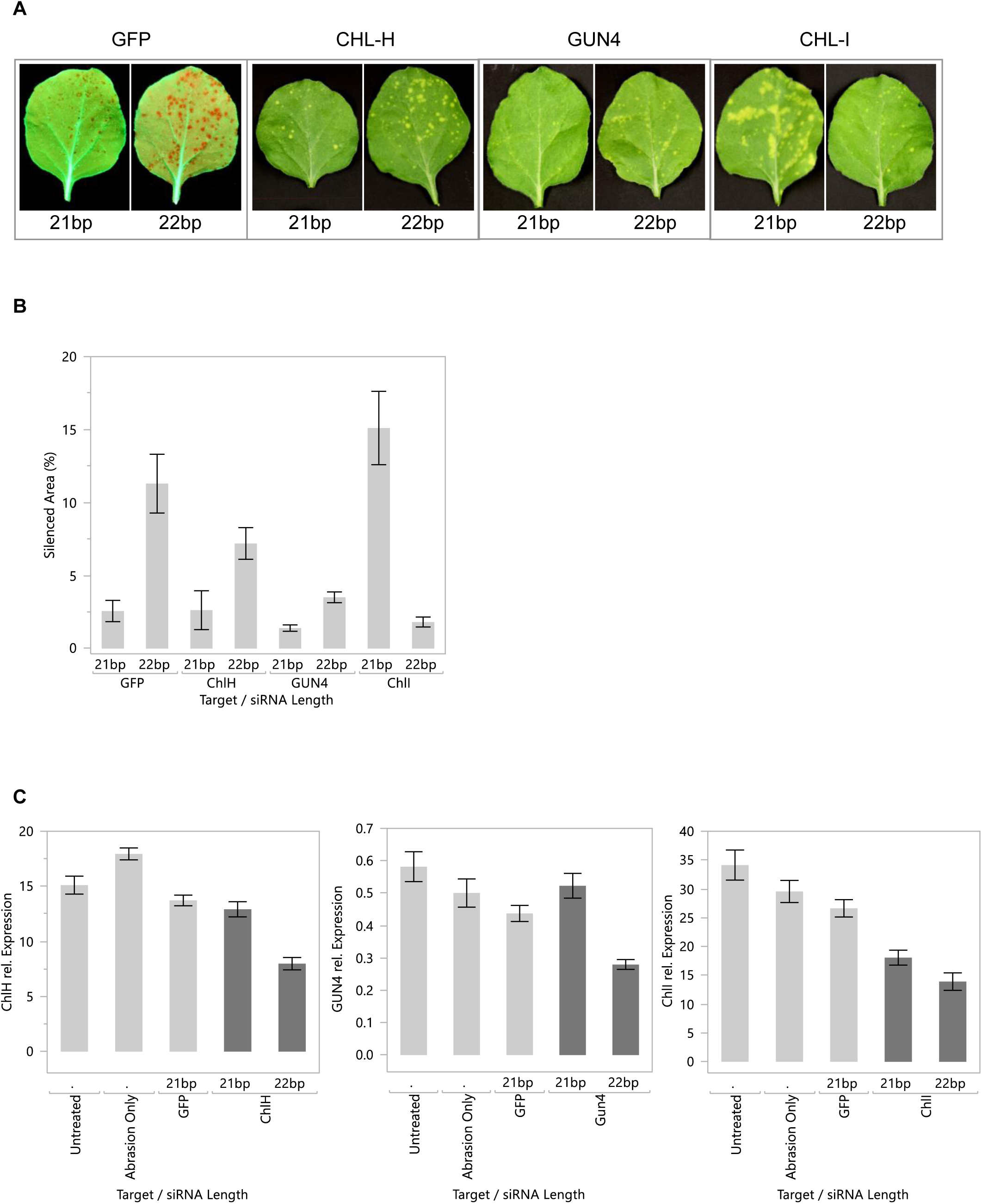
22 nt siRNAs often result in greater local silencing compared to 21 nt siRNAs. A, for each gene target, equal concentrations of 21 or 22 nt siRNAs were applied with sandpaper abrasion to the third leaf of 21-22 day old *N. benthamiana* plants. Plants were photographed under white (*CHL-I, CHL-H, GUN4*) or UV (*GFP*) light 7 days after treatment. N = 4 for GFP and N=6 for other gene targets. A leaf with a representative phenotype is shown. B, Silenced area for these treatments was quantified using Image J. C, Plants (8 per treatment) were treated as described above with siRNAs against endogenous genes, and phenotypic tissue was harvested 5 days later for RNA isolation and quantification of targeted mRNAs by RT-PCR. All data are means +/-standard error.

sRNA sequencing was carried out to examine the relative levels of transitive sRNAs produced in response to 22 nt siRNAs for these endogenous genes. siRNAs for *CHL-I, CHL-H* and *GUN4* (Tables 2 and 4; 1 μg/μl) were applied to the 2 youngest leaves of 3-week old *N. benthamiana* line 16C plants. Phenotypic tissue from the treated leaves was harvested 7 days after treatment. RNA was extracted from this tissue and processed for sRNA sequencing. The background levels of sRNAs for a given gene can be estimated by the number of transitive sRNAs produced in the absence of the targeting siRNA (Table 5). When targeting *CHL-H* with a 22 nt siRNA, a very high number of transitive sRNAs were produced (Table 5). As seen before (Table 3), only low levels of sRNAs were observed when targeting *CHL-I*. A low but significant number of transitive sRNAs were observed for the *GUN4* target (Table 5). In each of these cases, transitive sRNAs mapped 3’ to the applied siRNA binding site (data not shown). The ratio of transitive siRNAs for these genes after treatment with a 22 nt siRNA was approximately 32: 8: 1 (*CHL-H* > *GUN4* > *CHL-I*).

**Table 5.**
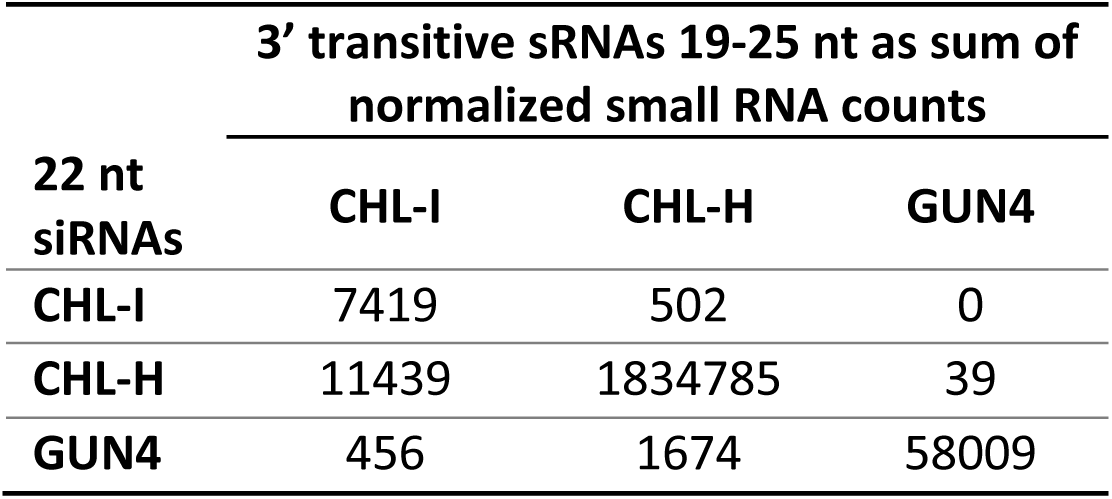
Sum of 3’ transitive small RNAs for endogenous genes targeted with 22 nt siRNAs.

| 22 nt siRNAs | 3' transitive sRNAs 19-25 nt as sum of normalized small RNA counts |  |  |
| --- | --- | --- | --- |
|  | CHL-I | CHL-H | GUN4 |
| CHL-I | 7419 | 502 | 0 |
| CHL-H | 11439 | 1834785 | 39 |
| GUN4 | 456 | 1674 | 58009 |

### Topical RNAi suppression of *CHL-H* gene in *Amaranthus cruentus*

To determine whether topically applied 22 nt siRNAs have similar properties in other species, we chose to target *CHL-H* in *Amaranthus cruentus*, a model for important weed species including Palmer Amaranth *(A. palmeri)* and Waterhemp *(A. tuberculatus)*. Similar to what was observed in *N. benthamiana*, a 22 nt siRNA was more effective in silencing *CHL-H* than was a 21 nt siRNA in *A. cruentus* with about 5 times more silenced area (Fig. 6A, B). The background levels of sRNAs for the CHL-H transcript in leaf tissue treated with a non-targeting dsRNA were very low, but significant numbers of transitive sRNAs were produced in response to 21 and 22 nt siRNAs, and these sRNAs mapped 3’ to the siRNA binding site (Table 6 and Fig. 6C). The level of transitive sRNAs produced in response to the 22 nt siRNA was about 12 times greater than those produced for a 21 nt siRNA. The sizes of transitive sRNAs for CHL-H in *A. cruentus* were mostly 21 nt with a significant number of 22 nt sRNAs also (Fig 6C insets). These results show that 21 and 22 nt siRNAs have similar effects in both *A. cruentus* and in *N. benthiamiana*. In both species, 22 nt siRNAs targeting CHL-H were more active than 21 nt siRNAs in producing a greater silencing phenotype and also by inducing a greater number of transitive sRNAs for many gene targets.

**Table 6.** Sum of sRNAs mapping to the CHL-H genes after treatment of *A. cruentus* with 21 or 22 nt siRNAs. Scr, scrambled dsRNA (Table 1).

| siRNA Type / Target | sRNA Type | Sum of normalized small RNA counts 19-25nt |
| --- | --- | --- |
| 21U / CHL-H | 5' Transitive | 5 |
| 21U / CHL-H | 3' Transitive | 1862 |
| 22U / CHL-H | 5' Transitive | 41 |
| 22U / CHL-H | 3' Transitive | 22679 |
| 22Con / Scr | background | 927 |

**Figure 6.**
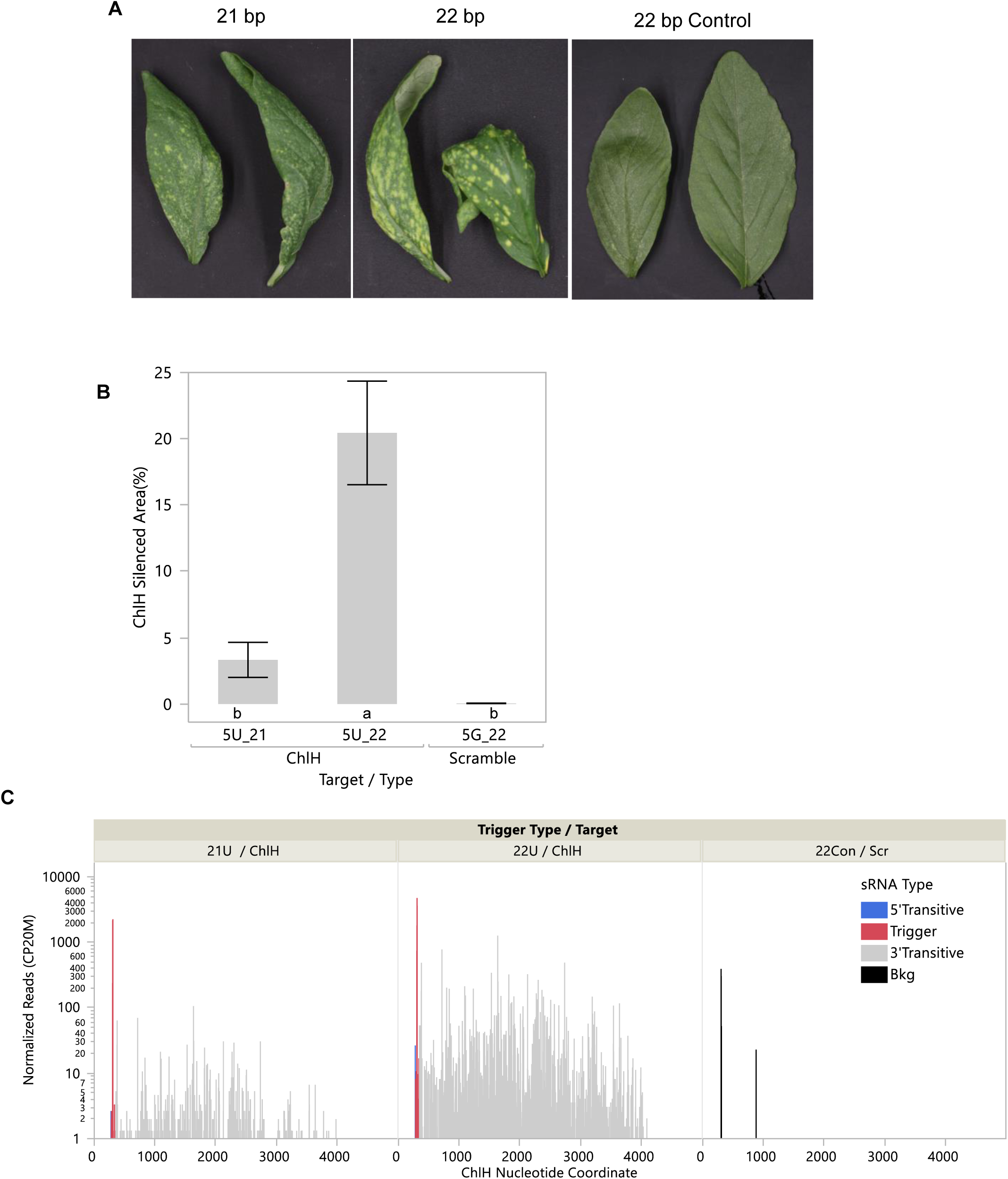
Topical treatment of *A. cruentus* leaves with siRNAs targeting *CHL-H* showed that 22-mers produced a stronger phenotype compared to 21-mers and resulted in more transitive sRNAs. siRNAs were applied to 14-day-old plants (6 reps), and treated leaves were abraded with dry particles. A, treated leaves were photographed 7 days after treatment, and leaves with representative phenotypes for the treatment are shown. B, Leaf silenced area was quantified using Image J, and plotted with standard errors. Letters under the bars indicate significant difference by Student’s t test, p<0.05. C, small RNAs (19-25 nt) were mapped to the targeted *CHL-H* gene.

## Discussion

We used a simple abrasion method for introduction of dsRNAs into leaf tissue and were able to demonstrate RNAi phenotypes for both the *GFP* transgene and 3 nuclear-encoded genes involved in chlorophyll biosynthesis or plastid signaling in *N. benthamiana* and for *CHL-H* in *A. cruentus*. We also examined production of secondary siRNAs in response to different dsRNA structures to explore the relationship between secondary siRNA production and systemic silencing.

We found that siRNAs from 20 to 22 bp were active in silencing a *GFP* transgene in the treated leaf tissue using this method (Fig 2). The requirement for a 22 nt RNA in the guide strand for induction of systemic silencing of the GFP transgene of *N. benthamiana* line 16C was confirmed using this method, and 22/22 and 21/22 nt siRNAs had equivalent efficacy (Fig 2). These results agree with reports using a high-pressure spray to deliver dsRNAs to silence GFP in the 16C line (Dalakouras et al. 2016). We also showed that a 22/21 structure, containing a 22 nt passenger strand, was not effective in generating a systemic response. Sequencing of sRNAs showed that a 22 nt RNA guide strand was important for the production of high levels of GFP transgene secondary siRNAs (Table 3 and Fig 4). However, a 22 nt RNA in the passenger strand did not induce transitive siRNA production. This result is in conflict with a previous report which found that having a 22 nt only in the passenger strand was sufficient to induce accumulation of secondary siRNAs (Manavella et al. 2012). We found good correspondence between the induction of high levels of secondary siRNAs and systemic silencing activity for the GFP gene in *N. benthamiana*.

To determine whether systemic silencing could be initiated for endogenous genes using topically applied dsRNAs, we targeted mRNAs for 2 sub-units of magnesium cheletase and the *GUN4* gene. Strong silencing phenotypes could be observed for each of these gene targets in the treated leaves (Figs 3B, 5, 6) but no systemic silencing was observed for any of them. sRNA sequencing showed that 22 nt siRNAs induced some transitive sRNA production for each of these endogenous genes (Figs 4 and 6, Tables 5 and 6). However, the amount of secondary sRNAs was quite different for the different gene targets. Only a very low level of secondary siRNAs was produced in response to a 22 nt siRNA for *CHL-I* (Tables 3 and 5), and this response was only observed with a 22 nt siRNA and not with a 21 nt siRNA. Somewhat higher levels of secondary siRNAs were produced for the *GUN4* mRNA after treatment with a 22 nt siRNA, but very high levels of secondary siRNAs were produced for the *CHL-H* mRNA in *N. benthamiana* (Table 5). Similarly, high levels of secondary siRNAs were also observed for *CHL-H* in *A. cruentus* after targeting with a 22nt siRNA (Table 6). Despite these high levels of secondary siRNAs for *CHL-H*, no systemic silencing was observed in either species. In subsequent experiments, greater levels of local silencing for *CHL-H* were achieved, and yet systemic silencing was still not observed (data not shown). Therefore, elevated amounts of secondary siRNAs were not sufficient for induction of systemic silencing. Since we didn’t compare secondary siRNAs for *GFP* and *CHL-H* in the same experiment, it’s not possible to address whether a threshold of transitive sRNA concentration exists that is required to initiate systemic silencing. Published reports of systemic silencing have focused on examples using transgenes, either stably integrated or transiently expressed. In all of these cases very high levels of target and siRNAs or amiRNAs would be present.

The distribution of secondary siRNAs along the coding sequence were different for the *GFP* transgene compared to the endogenous genes examined in this study. For *GFP*, secondary siRNAs were produced both 3’ and 5’ to the targeting siRNA (Fig 4A and Table 3). However, for *CHL-I* (Fig 4B and Table 3) and for *CHL-H* (Fig 6 and Table 6) secondary siRNAs were mostly observed 3’ to the targeting siRNA site. It’s possible that presence of secondary siRNAs 5’ to the targeting siRNA could be associated with systemic silencing, but additional examples are needed.

Although production of secondary siRNAs with 22 nt siRNAs didn’t result in systemic silencing of endogenous genes, an increase in the amount of local silencing was observed for both *CHL-H* and *GUN4* targets (Figs. 5 and 6). In contrast, for the *CHL-I* gene silencing was greater with the 21 bp siRNA. It is possible that the 22 bp siRNA had lower silencing efficacy for *CHL-I* due to the very weak induction of secondary sRNAs by 22 nt siRNAs for this gene target (Tables 3 and 5). This weak induction of secondary small RNAs may be an intrinsic property of the *CHL-I* gene.

The sandpaper abrasion method for introduction of sRNAs into plant cells is easy to use, highly reproducible, and results in RNAi silencing of the targeted genes. Although plant tissues are known to contain abundant RNAses in both the apoplast and within the cell (Green 1994; Pérez-Amador et al. 2000), we found specific effects of 21 vs 22 nt siRNAs, suggesting that at least some of the siRNAs are preserved intact during uptake into cells. This method can be used for additional investigation of RNAi mechanisms in plant cells.

## Materials and Methods

### Plant Materials

The *N. benthamiana* line 16C expressing a *GFP* transgene (Ruiz et al. 1998) was obtained from David Baulcombe (Ruiz et al. 1998). Plants were grown in a growth chamber under a 16 h day: 8 h night photoperiod. Light intensity was approximately 250 μmol m^-2^ s^-1^ and temperatures were 26°C during the day and 18°C at night.

*Amaranthus cruentus* seed was obtained from Johnny’s Selected Seed (https://www.johnnyseeds.com/). Plants were grown at 25°C with 16 h day length and 100 μmol m^-2^ s^-1^ light intensity. Seeds were germinated on coconut coir plugs (Jiffy Products of America, Inc., Lorain, OH) and then transplanted on day 7 into 2.5in pots filled with Berger BM2 media. Plants were irrigated daily with an ebb and flow system with Peters 20-20-20 liquid fertilizer.

### Topical application of dsRNAs to plant leaves

siRNAs (20-22 bp) were synthesized by IDT (Integrated DNA Technologies, Inc., Iowa, USA).

#### N. benthamiana

Typically, 20-25 μl siRNA at 1 μg/μl in 0.01%-0.05% Silwet L77 or in water was spread onto 2-3 partially expanded leaves of 14 to 21-day-old *N. benthamiana* plants using a pipet tip. Sandpaper (#600 grit) was gently rolled onto the treated leaf surface using support from a gloved finger, taking care not to tear the leaf.

For the experiment shown in Figure 5, the concentration of each 22 nt siRNA that allowed resolution of individual phenotypic spots was determined empirically. Then, equal concentrations of 21 and 22 nt siRNAs for a gene target were compared for activity. siRNA concentrations used were: CHL-H (0.5 μg/μl); GUN4 (0.5 μg/μl); GFP (1 μg/μl); CHL-I (0.2 μg/μl).

#### Amaranthus cruentus

dsRNA was dissolved at 3 mg/ml (21/22 mers) or 6 mg/ml (66 mer) in 50mM Sucrose, 5mM MES pH 5.7, 0.08% L77 Silwet. dsRNA solution (8μl) was applied to each of the L3, L4 and terminal leaves and allowed to air dry. The leaves were then abraded with 220 mesh silicon carbide particles (Kramer Industries) using a custom-built dry particle sprayer equipped with a 95015-TC spray tip (Spraying Systems Co.) at a pressure of 45 psi, a speed of 36.5cm/s, set 15 cm from the leaf surface.

### Phenotype image capture and analysis

For young plants GFP fluorescence was excited using a high intensity blue LED light source (465 nm, Photon System Instruments) and imaged using a Cannon EOS 70D camera with 58 mm Tiffen Green #11 and Yellow #12 filters. GFP fluorescence of older plants was visualized using a UV light and images captured using an iPhone 7.

Phenotypes were quantified using the Image J 1.51a platform (National Institutes of Health, USA). The threshold color panel was used to distinguish phenotypic regions of leaves. Both the leaf border and the phenotypic borders were manually outlined. Then, the program automatically quantitated the pixel number within the whole leaf or the phenotypic region, and silenced area was represented by the ratio of the phenotypic area pixels to the total leaf area pixels. Color threshold settings were kept uniform within experiments.

Photographs used for figures were cropped and adjusted for brightness, contrast, hue and saturation using Paint.net. All images from an experiment were adjusted using the same settings.

### Sample collection and RNA purification

Leaf discs were collected from treated leaves using 5 mm biopsy punches (INTEGRA YORK PA INC.) and were immediately frozen on dry ice and stored at −80°C. Total RNAs were extracted from 10 disks per sample using Trizol reagent (Invitrogen, CA, USA) following the instructions of the product manual.

### RNA gel blot

The RNA gel blot was done by using the DIG Northern Kit (Roche, Mannheim, Germany). The blot was probed with part of the *GFP* transcript (Supplementary Table 1).

### RNA Detection by RT-qPCR

Frozen leaf disc samples were ground with ball bearings and extracted using Direct-zol™-96 RNA (Zymo Research; Irvine, CA). Total RNA was eluted in DNAse/RNAse free water, quantified to ensure the concentration was less than 200ng/ul and used directly for cDNA synthesis. Oligo dT cDNA was generated with High Capacity cDNA Reverse Transcription Kit (Applied Biosystems; Foster City, CA) per the manufacturer’s protocol. Singleplex qPCR was performed with a 4-fold dilution of cDNA using either Quanta Bio (Beverly, MA) PerfeCTa Fast Mix II for probe based assays or PerfeCTa SYBR green Supermix and run on BioRad CFX96 Real-Time PCR Detection System (Hercules,CA). For quantification, EF1a and UKN1 were used as internal normalizer genes. Relative Quantity (RQ) was calculated by 2^-(GOI Ct – GeoMean Internal Normalizer Ct). Probe sequences are shown in Supplementary Table 2.

### sRNA sequencing and analysis

Illumina’s TruSeq small RNA Library Preparation Kit was used to prepare small RNA libraries following the manufacturer’s protocol (Document # 15004197v02) with slight changes to the library purification steps. After cDNA amplification, uniquely indexed libraries were pooled together in equimolar amounts and size separated on a 6% Novex TBE PAGE Gel. The gel was stained for twenty-minutes with 1X SYBR Gold. A minimum of 3 million reads per size-selected library was generated using Illumina’s NextSeq platform.

To assess quality of the sequencing libraries, fastqc was used (Andrews, 2010, http://www.bioinformatics.babraham.ac.uk/projects/fastqc), and for read-quality filtering and trimming adapters, Trimmomatic was used (Bolger et al. 2014). Read processing and mapping was performed with SAMtools (Li et al. 2009), BAMtools (Barnett et al. 2011), and bowtie2 using very-sensitive-local alignment parameters (Langmead and Salzberg 2012), with further analysis done using custom shell and R scripts. To facilitate comparisons among libraries, raw counts of mapped reads were normalized to the total number of reads passing both length (18-48 nt) and quality criteria (5 base sliding window with average quality above 20).

## Acknowledgments

We thank Professor David Baulcombe for sharing the *N. benthamiana* 16C *GFP* line. We acknowledge excellent plant production by Brenda Reed and Kaylene Yandel, and seed production by Jim Byrne. Our image analysis protocol was developed by Andrew Mroczka. We thank the Bayer Sequencing team for generation of sRNA libraries and sequencing.

---

**Table S1.**
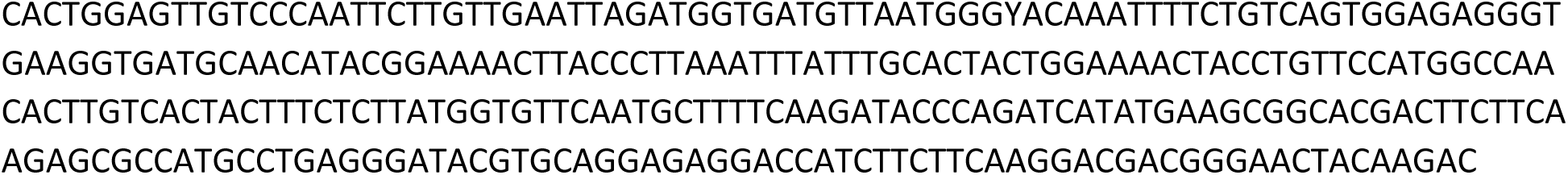
*GFP* 5’ probe sequence for RNA gel blot.

**Table S2.**
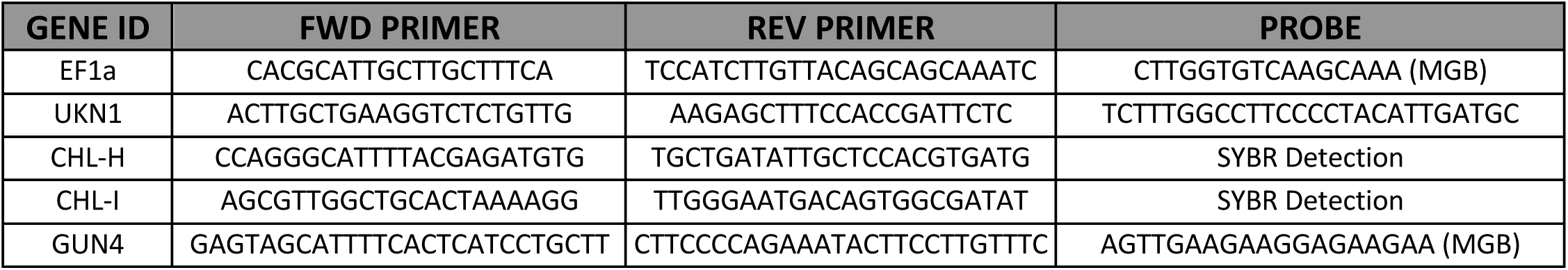
RT-qPCR primer and probe sequences for transcript measurements (Sequences are 5’-3’). MGB, Minor groove binding probe.

